# Human rhinovirus 16 impairs macrophage cytokine secretion by disrupting NF-κB nuclear translocation and intracellular cytokine trafficking

**DOI:** 10.64898/2026.07.23.740013

**Authors:** Zoé Fremont-Debaene, Zeyni Mansuroglu, Léane Puchot, Marjorie Leduc, Frédéric Bonhomme, Paola B. Arimondo, Florence Niedergang, Suzanne Faure-Dupuy

## Abstract

Human rhinovirus (HRV) infections are a major cause of acute exacerbations in chronic obstructive pulmonary disease (COPD), often promoting secondary bacterial infections by dysregulating macrophage function. Although HRV16 has previously been shown to impair macrophage cytokine secretion, the underlying molecular mechanisms remain poorly understood.

To address this, we examined the effects of HRV16 on primary human monocyte-derived macrophages, subsequently challenged with lipopolysaccharide (LPS) to mimic secondary bacterial infection. HRV16 significantly reduced IL-10 and IL-1β expression at both the mRNA and protein levels. In contrast, IL-6 transcription was increased despite markedly reduced cytokine secretion. Immunofluorescence analysis revealed enhanced colocalization of IL-6 with the Golgi apparatus following HRV16 infection, consistent with intracellular retention and defective trafficking. These findings reveal that HRV16 disrupts cytokine secretion through distinct transcriptional and post-transcriptional mechanisms.

To investigate the basis of transcriptional dysregulation, we performed quantitative histone post-translational modification profiling by mass spectrometry, which identified multiple HRV16-induced epigenetic alterations. Notably, a decrease in the active epigenetic mark H2AZK4Ac was observed. Chromatin immunoprecipitation demonstrated unchanged H2AZK4Ac occupancy at the IL-10 and IL-1β promoters but increased enrichment at the IL-6 promoter, consistent with its selective transcriptional upregulation. HRV16 infection also induced sustained phosphorylation of NF-κB p65 that was accompanied by impaired nuclear translocation, suggesting defective activation of NF-κB-dependent transcription.

Together, these results demonstrate that HRV16 inhibits cytokine secretion through disruption of NF-κB signalling and defective intracellular cytokine trafficking and identify associated alterations in the macrophage epigenetic landscape. These findings provide new mechanistic insight into rhinovirus-mediated dysregulation of macrophage inflammatory responses and its potential contribution to impaired antibacterial immunity during COPD exacerbations.

## Introduction

Macrophages are essential components of the innate immune system and constitute the first line of defense against invading pathogens^1^. As highly plastic immune cells, macrophages play central roles in pathogen sensing^2^, phagocytosis^3^, antigen presentation^4^, tissue remodeling^5,6^, and the orchestration of inflammatory responses via cytokine secretion^1^.

Pathogen recognition by macrophages is largely mediated by pattern recognition receptors (PRRs), including Toll-like receptors (TLRs)^7,8^, which detect conserved pathogen- associated molecular patterns (PAMPs). Upon activation, TLRs initiate intracellular signaling cascades that trigger the activation of transcription factors such as nuclear factor kappa B (NF-κB), interferon regulatory factors (IRFs), and activator protein 1 (AP-1), thereby promoting the expression of inflammatory cytokines, chemokines, and antiviral mediators^7^. TLR signaling pathways are tightly regulated and can differ depending on receptor localization and adaptor recruitment. Surface TLRs, such as TLR4, primarily detect bacterial products (e.g., lipopolysaccharide or LPS)^9^, whereas endosomal TLRs, such as TLR3^10^, recognize viral nucleic acids.

TLR4 signaling is particularly complex because it activates both MyD88-dependent and TRIF-dependent pathways^9^. The MyD88-dependent pathway rapidly induces NF-κB activation via the degradation of its inhibitor IκBα, which must then be resynthesized to induce a negative feedback loop^11^. NF-κB activation drives the transcription of pro- inflammatory cytokines such as interleukin1 beta (IL-1β), IL-6 and tumor necrosis factor alpha (TNFα) as well as anti-inflammatory cytokines such as IL-10^11^. In parallel, the TRIF- dependent pathway activates IRF3 and contributes to the expression of interferon- stimulated genes and antiviral responses^9^. In contrast, TLR3 signals mainly through TRIF^12^.

In addition, gene expression can also be regulated by epigenetic modifications such as DNA methylation and histone modifications that dynamically regulate chromatin accessibility and transcription factor recruitment^13^. Histone acetylation is generally associated with transcriptionally active chromatin, histone methylation can either activate or repress transcription depending on the modified residue, whereas DNA methylation represses transcription^13^. During infection, innate immune cell can undergo profound chromatin remodeling that shapes cytokine production and inflammatory responses^13,14^.

Several studies have shown that viruses can manipulate macrophages-mediated cytokine secretion to suppress antiviral response, thereby facilitating immune evasion and viral persistence^15,16^.

The human rhinovirus 16 (HRV16), a member of the Enterovirus genus within the Picornaviridae family, is one of the principal causative agents of the common cold^17^. Although HRV infections are generally mild and self-limiting in healthy individuals, they represent a major health burden in patients suffering from chronic respiratory diseases such as asthma and chronic obstructive pulmonary disease (COPD)^18,19^. HRVs are among the viruses most frequently detected during COPD exacerbations, which are often associated with secondary bacterial infections and severe inflammatory responses^20,21^. These exacerbations significantly contribute to COPD-related morbidity and mortality worldwide.

Accumulating evidence indicates that HRV16 profoundly alters macrophage functions usually involved in antibacterial immunity. We have shown that HRV16 impairs bacterial internalization^22^, phagosome maturation and bacteria killing ^23,24^, and cytokine secretion^25^ in macrophages, thereby creating a permissive environment for bacterial superinfections. However, the molecular mechanisms underlying HRV16-mediated cytokine secretion inhibition remain to be deciphered.

In this study, we sought to elucidate the molecular mechanisms by which HRV16 impairs macrophage inflammatory responses. Using primary human monocyte-derived macrophages, we investigated the effects of HRV16 exposure on TLR signaling pathways, epigenetic regulation, and cytokine secretion mechanisms. We demonstrate that HRV16 impairs NF-κB p65/RelA nuclear translocation while preserving IRF3 signaling, resulting in defective transcription of specific inflammatory cytokines. In parallel, HRV16 disrupts intracellular trafficking, leading to impaired cytokine secretion.

Together, our findings reveal that HRV16 alters macrophage inflammatory responses through both transcriptional and post-transcriptional mechanisms, providing new insights into viral immune evasion strategies that may contribute to bacterial superinfections.

## Materials and Methods

### Monocytes-derived macrophages purification and differentiation

Human Monocytes-derived Macrophages (hMDMs) were obtained from the purification of peripheral blood mononuclear cells (PBMCs) isolated from whole blood. Whole blood is collected from healthy donors at the Etablissement Franc ais du Sang (INSERM agreement #18/EFS/030 and CNRS agreement #2023-2026-025 CCPSL CNRS) ensuring that all donors gave a written informed consent and providing anonymized samples. The procedures for sample collection and conservation were declared and approved through CODECOH (COnservation d’Eléments du COrps Humain, No. DC-2021-4166) by the French Ministry of Higher Education, Research and Innovation, in accordance with French regulations on human biological elements. PBMCs were isolated by density gradient (Ficoll-Plaque Plus, Cytiva, ref.17144003) and monocytes were further isolated by a Percoll (VWR, ref. 17-0891) density gradient. Monocytes were plated in plastic plates (24 or 12 well plates, 2.5 ×10^5^ cells per cm^2^) or 15 cm culture dish (1×10^5^ cells per cm^2^) with 4 ng/mL human GM-CSF (Miltenyi, ref. 130-093-865) and 0.5 ng/mL M-CSF (Miltenyi, ref. 130-093-963) in R10 medium (RPMI 1640 (Gibco, ref. 11875093) + 10% Fetal Bovine Serum (FBS; Gibco, ref. 102701006) + 100 µg/ml streptomycin/penicillin (Sigma-Aldrich, ref. P4333) + 10 mM HEPES (Gibco, ref. 15630080)+ 1mM Sodium Pyruvate (Gibco, ref. 2009465) + 1X non-essential amino acids (Gibco, ref. 11140050)). After 4 days, the medium was replaced by fresh R10 medium supplemented with cytokines. After 7 days, the medium was changed to MONO medium (RPMI 1640 + 10% Fetal Bovine Serum + 2 mM L-glutamine + 100 µg/ml streptomycin/penicillin). Cells were cultured at 37°C with 5% CO2. After 10 days of differentiation, primary hMDMs were used as described in the figures’ legends.

### HeLa Ohio culture

HeLa Ohio cells were purchased from the European Collection of Authenticated Cell Cultures (ECACC, ref. 84121901) and were cultured at 37 °C with 5% CO2 in HeLa Ohio medium (DMEM GlutaMax containing 25 mM D-glucose (Gibco, ref. 10569010) supplemented with 10% Fetal Bovine Serum + 2 mM L-glutamine + 100 µg/ml streptomycin/penicillin). Cells were cultured at 37°C with 5% CO2. The cells were passaged every 3 days.

### HRV16 production

Human rhinovirus 16 (HRV16) (VR-283, strain 11757, lot 62342987) was purchased from the ATCC and stocks were produced by infecting HeLa Ohio cells in virus medium (DMEM GlutaMax containing 25 mM D-glucose supplemented with 10% FBS, and 2 mM L-glutamine). HeLa Ohio cells were plated in 6 well plates and grown to 80% confluence at 37 °C with 5% CO2 and infected with 300 μL/well of HRV16 at 0.25 × 10^7^ TCID50/mL or control media (Mock) for 1h at room temperature with agitation. Then the medium was completed to a final volume of 2 mL with virus media. After that, the cells were left for 48h at 37°C with 5% CO2. When the cells reached 90% of infected cells (cells get round), the plates (cells + supernatants) were frozen at -20°C and unfrozen a total of 3 times to lyse the cells. Finally, the supernatants were centrifuged at 4,000 rpm for 15 min, filtered at 0.22 μm, and stocks were stored at −80 °C.

### Quantification of the viral TCID50

HeLa Ohio cells were cultivated in a 96-well plate to reach 80% confluence. HRV16 or Mock stocks were diluted from 1:10^0^ to 1:10^11^ dilution. Dilutions were done using the virus medium. The different dilutions were then added to the cells for a total of 6 replicates for each dilution for the virus and 2 replicates for each dilution of the Mock. Cells were then incubated for 4 days at 37°C with 5% CO2. Wells showing cell lysis were counted and TCID50 was calculated using the Spearman-Karber formula.

### Viral exposure and stimulation of hMDMs

Cells were exposed to 70 μL for 24-wells plates, 150 μL for 12-wells plates, 300 μL for 6- wells plates and 5 mL for 15 cm culture dishes of HRV16 at a concentration of 1 × 10^7^ TCID50/mL for 1h at room temperature under agitation. As controls, cells were exposed to the same amount of Mock stock. Then the cells were washed with phosphate-buffered saline (PBS; Gibco, ref. 14190-094). After at the indicated time after infection (see figures’ legends), cells were stimulated with TLR ligands for TLR3 and TLR4. TLR ligands used for the stimulation are: LPS-B5 (Invivogen, ref. tlrl-b5lps) at 100ng/mL for TLR4 and Poly(I:C) High Molecular Weight (Invivogen, ref. tlrl-pic) at 5μg/mL for TLR3. Cells were stimulated for the indicated amount of time (see figures’ legends).

### RNA extraction, reverse transcription, and qPCR

The cells were lysed in 350 μL of LBP buffer (Macherey-Nagel, ref. 740984.250) at either 0h, 4h, 8h, 24h, 48h or 72h post HRV16 exposure (hpe). Alternatively, the cells were stimulated at either 24hpe, 48hpe, 72hpe, 96hpe or 120hpe and lysed 6h post stimulation.

RNA was extracted using the NucleoSpin RNA Plus kit (Macherey-Nagel, ref. 740984.250) according to the manufacturer’s instructions. Reverse Transcription was done using the High-Capacity cDNA Reverse Transcription Kit (Thermo Fisher, ref. 4368813), using 250 ng of RNA and following manufacturer’s instructions. The cDNA was then diluted to a final concentration of 2.5 ng/mL in RNAse free water.

qPCR was performed using the SYBR Green I Master (Roche, ref. 04707516001) with specific oligos **(Supplementary Table 1)**. For the qPCR mix, 1 μL of diluted cDNA, 0.2 μL of the corresponding forward/reverse primers at a concentration of 10 μM, 5 μL of SYBER Green, and 3.6 μL of RNAse free water were used. The qPCR was performed on a LightCYler480 (Roche) with the following program: 95 °C for 2min, then 45 amplification cycles at 95 °C for 5s, 60 °C for 10s, and 72 °C for 15s, followed by a dissociation step at 95 °C for 5s and then 65 °C for 1min. TBP (TaTa-box Binding Protein) was used as a reference gene. The data analysis was done using the ΔΔCT method, for which samples were normalized either to the average of the Mock at 24hpe in the experiment without any TLR4 stimulation or to the average of the Mock+TLR4 stimulation at 24hpe for the other experiments (see Figures’ legends).

### ELISA

The cells were stimulated with TLR4 ligand either at 24hpe or 48hpe and the supernatants were recovered 6h or 24h post stimulation. The ELISA were done using the Bio Techne DuoSet ELISA kits to detect IL-1β (ref. DY201-05), IL-6 (ref. DY206-05) and IL- 10 (ref. DY217B-05) following the manufacturer’s protocol. Briefly: the plates (Thermoscientific Nunc MaxiSorp, ref. 442404) were coated overnight with the capture antibody diluted in PBS. The following day, plates were saturated with PBS + 1% bovine serum albumin (BSA; Euromedex, ref. 04-100-812E) for 1h. Samples diluted or not in PBS + 1% BSA and standards were added to the plate (Pure samples for IL-1β, 1:2 dilution for IL-10, and 1:10 dilution for IL-6). After 2h of incubation, the detection antibody diluted in PBS + 1%BSA was added and incubated for 2h. Then the streptavidin-HRP diluted in PBS + 1%BSA was added for 20 min away from light then removed. The results were revealed by adding TMB (Sigma-Aldrich, ref. T0440) and the reaction was stopped by adding a stop solution (2N H2SO4). Between each step, the plates were washed repeatedly. Then, the plates were read with a Multiskan FC (ThermoFisher). The results are expressed as concentration of the indicated cytokine in pg/mL.

### DNA extraction and DNA methylation analysis

At 24hpe, the cells were stimulated or not with TLR4 ligand for another 24h. Then DNA was extracted using the Qiagen DNAeasy blood and tissue kit (ref. 69504) according to the manufacturer’s instructions. DNA was digested using the Nucleoside Digestion Mix kit (New Englands Biolabs, ref. M0649S). Then, analysis of global levels of 5mC and 5hmC were performed on a Q exactive mass spectrometer (Thermo Fisher Scientific). It was equipped with an electrospray ionization source (H-ESI II Probe) coupled with an Ultimate 3000 RS HPLC (Thermo Fisher Scientific).

Digested DNA was injected onto a ThermoFisher Hypersil Gold aQ chromatography column (100 mm * 2.1 mm, 1.9 μm particle size) heated at 30°C. The flow rate was set at 0.3 mL/min and run with an isocratic eluent of 1% acetonitrile in water with 0.1% formic acid for 10 minutes.

Parent ions were fragmented in positive ion mode with 10% normalized collision energy in parallel-reaction monitoring (PRM) mode. MS2 resolution was 17,500 with an AGC target of 2e5, a maximum injection time of 50ms, and an isolation window of 1.0 m/z.

The inclusion list contained the following masses: C (228.1), 5mC (242.1), and 5hmC (258.1). Extracted ion chromatograms of base fragments (±5ppm) were used for detection and quantification (112.0506 for C; 126.0662 for 5mC; 142.0609 for 5hmC). Calibration curves were previously generated using synthetic standards (CliniSciences, France) in the ranges of 0.2 to 40 pmol injected for C and A and 0.01 to 1 pmol for 5mC and 5hmC. Results are expressed as a percentage of total C or A and normalized for each donor on the Mock-non stimulated condition.

### Histone purification and mass spectrometry analysis of epigenetic marks on histones

At 24hpe, the cells were stimulated with TLR4 ligand for 24h, as described above. On the first day of the purification, cells were scrapped in PBS and centrifuged 10min at 4°C and at 3,500 rpm. The pellet was then resuspended in a NB1 buffer composed of CaCl2 5 mM, MgCl2 5 mM, KCl 60 mM, Saccharose 250 mM, NaCl 15 mM, Tris 15 mM, DTT 1 mM and 1 protease inhibitor cOmplete^TM^ EDTA-free tab (Roche, ref. 11873580001). Then, resuspended cells were centrifuged 10min at 4°C and at 1,000 g. The pellet was then resuspended in a NIB2 buffer composed of NIB1 buffer + NP40 0,2% to lyse the plasmic membrane. After 10min of incubation, the lysate was centrifugated 10min at 4°C and at 1,000 g. Then the pellet was washed 3 times by resuspending in NIB1 followed by centrifugation at 10min at 4°C and at 1,000 g. After the washes, the pellet was resuspended in 0.2 M of H2SO4 and incubated 4h at 4°C. After a centrifugation of 10min at 4°C and at 5,000 g, the supernatant was recovered and mixed with Trichloroacetic acid (TCA) 100% and incubated at 4°C overnight. On the second day of purification, the supernatant + TCA was centrifugated 10min at 4°C and at 5,000 g to pellet the histones. The histones were then washed 2 times by resuspending it with acetone -20° and then a 10min at 4°C and at 5,000 g centrifugation. Finally, the histones were resuspended in acetic acid, and the purification was tested with a SDS-PAGE and Coomassie Blue coloration. Then the samples were sent to the PROTEOM’IC facility of Institut Cochin where sample were processed as follow:

#### Samples digestion

150 µl of purified histones samples were separated to 3 tubes (HL,HT and HpT) for guanidium chloride digestions. To do so, they were dried using a speedvac. The HpT tubes were resuspended with triethylamonium bicarbonate 0.5 M and propionylated for 15min at room temperature with 40 µl of 75% acetonitrile, 25% anhydride propionic. These samples were then dried. HL, HT and HpT samples were resuspended with 40 µl of tris(2- carboxyethyl) phosphine 10 mM, chloroacetamide 40 mM, guanidium chloride 6 M and Tris 100 mM pH8.5 and heat for 5min at 95°C. Samples were diluted with 400µl of Tris25mM pH8.5, acetonitrile 10%. 0.3 µg of trypsine were added to HT and HpT samples. 0.3 µg of LysC were added to HL samples. All samples were incubated at 37°C overnight for digestion. Peptides samples were purified using C18 stagetip and dried.

#### LC-MS/MS

Peptides samples were resuspended with 10 µl of acetonitrile 10%, trifluoroacetic acid 0.1%. LC-MS analyses were performed on a Dionex U3000 HPLC nanoflow chromatographic system (Thermo Fischer Scientific, Les Ulis, France) coupled to a TIMS- TOF HT mass spectrometer (Bruker Daltonik GmbH, Bremen, Germany). 1 μL was loaded, concentrated and washed for 3min on a C_18_ reverse phase precolumn (3 μm particle size, 100 Å pore size, 75 μm inner diameter, 2 cm length, from Thermo Fisher Scientific). Peptides were separated on an Aurora C18 reverse phase resin (1.6 μm particle size, 100Å pore size, 75 μm inner diameter, 25 cm length mounted onto the Captive nanoSpray Ionisation module, (IonOpticks, Middle Camberwell Australia ) with a 1h overall run-time gradient ranging from 99% of solvent A containing 0.1% formic acid in milliQ-grade H2O to 55% of solvent B containing 80% acetonitrile, 0.085% formic acid in mQH2O. The mass spectrometer acquired data throughout the elution process and operated in DDA PASEF mode with a 1.1 second/cycle, with Timed Ion Mobility Spectrometry (TIMS) enabled and a data-dependent scheme with full MS scans in Parallel Accumulation and Serial Fragmentation (PASEF). This enabled a recurrent loop analysis of most intense nLC-eluting peptides which were CID-fragmented between each full scan every 1.89 second. Ion accumulation and ramp time in the dual TIMS analyzer were set to 166 ms each and the ion mobility range was set from 1/K0 = 0.6 Vs cm^−2^ to 1.6 Vs cm^−2^. Precursor ions for MS/MS analysis were isolated in positive polarity with PASEF in the 100-1,700 m/z range by synchronizing quadrupole switching events with the precursor elution profile from the TIMS device. Singly charged precursor ions were excluded from the TIMS stage by tuning the TIMS using the otof control software, (Bruker Daltonik GmbH). Precursors for MS/MS were picked from an intensity threshold of 1,000 arbitrary units (a.u.) and re-fragmented and summed until reaching a ‘target value’ of 20,000 a.u., while allowing a dynamic exclusion of 0.40 s elution gap.

#### Data analysis

The mass spectrometry data were analyzed using Maxquant version 1.6.17.0^26^. The database used was a Human histones sequences from El Kennani et al 2017^27^ and the list of contaminant sequences from Maxquant. Cystein carbamidomethylation was set as constant modification for all samples and propionylation of lysine was set as constant for HpT samples. Oxidation of methionine, Acetylation of Lysines, Trimethylation of lysines, Dimethylation of lysines and ariginines, Methylation of lysines and arginines were set as variable modifications for all samples. Second peptide search and the “match between runs” (MBR) options were allowed. False discovery rate (FDR) was kept below 1% on both peptides and proteins.

#### Statistical analysis

Modified sites quantifications were analysed using homemade shiny script: Proxima (https://github.com/MarjorieLeduc). For each digestion (HL, HT, HpT) of each modified site, intensities of HRV16 were compared to Mock ones by a welch test. P-value threshold was set to 0.05 and appeared/disappeared modified sites were also searched as sites with no intensity in one condition and 3 to 5 intensities in the other.

### Chromatin Immunoprecipitation (ChIP)

At 24hpe, the cells were stimulated with TLR4 ligand for 6h. Cells were washed in PBS and incubated for 10min with 1% formaldehyde at room temperature under agitation to crosslink DNA and proteins. Formaldehyde was removed and crosslinking was stopped with a 5min incubation with 0.125 M glycine in PBS at room temperature under agitation. The cells were then washed once in PBS and scrapped in 1mL of PBS. Cells were centrifuged 10min at 4°C and at 3,500 rpm. Pellets were washed once in PBS + 1 mM PMSF (phenylmethylsulfonyl fluoride, UGA, ref. 10485015) + 1X PIC (Proteinase inhibitor cocktail) (Roche, ref. 11836145001) and centrifugated again at 10min at 4°C and 3,500 rpm. The pellets were then recovered, resuspended in lysis Buffer (50 mM Tris HCl pH 8, 10 mM EDTA, 1% SDS, 1X PIC, 1 mM PMSF) and sonicated (6 cycles of 30s ON/30s OFF, high frequency) using a BioruptorPico (Diagenode, ref. B01060010). Chromatin fragments were recovered by centrifugation (10min at 4°C and 13,000 rpm) and diluted 9 times with Ripa Buffer (10 mM Tris HCl pH 7.5, 140 mM NaCl, 1 mM EDTA, 0.5 mM EGTA, 1% Triton, 0.1% SDS, 0.1% Na-Deoxycholate, 1X PIC, 1 mM PMSF). The chromatin was then precleared by incubation with 20 μL of Protein A-Sepharose beads (Millipore, ref. 16157) on a spinning wheel at 4°C for 2h. Samples were centrifuge for 1min at 4°C and 1,200 rpm to pellet Protein A-Sepharose beads. Chromatin was collected and incubated with the 6 μg of the corresponding antibody overnight at 4°C on the spinning wheel. The antibodies used for this experiment are: H3K27Me3 rabbit mAb (Ozyme, ref. C36B11), H3K27Ac rabbit mAB (Diagenod, ref. C15410196), H2AZK4+7+11Ac rabbit pAb (Abcam, ref.232908), and normal rabbit IgG (negative control; Cell signaling, ref. 2729s). The following day, the chromatin + antibodies complexes were immunoprecipitated by 2h incubation at 4°C on a spinning wheel with magnetic beads (Invitrogen, ref. 1002D) previously washed with Ripa Buffer. Then, using a magnet, the supernatants of the negative control conditions were recovered to serve as chromatin input. The beads were washed 4 times with Ripa Buffer, 1 time with LiCL Buffer (0.25 M LiCl, 0.5% NP40, 0.5% Na-Deoxycholate, 10 mM Tris HCl pH 8, 1 mM EDTA, 1X PIC, 1 mM PMSF), and 1 time with TE Buffer (10 mM Tris HCl pH 8, 10 mM EDTA). Sample were transferred in a new Eppendorf to reduce background. To reverse the crosslinking, the beads were incubated in 150 μL of elution buffer (20 mM Tris HCl pH 7.5, 5 mM EDTA, 50 mM NaCl, 1% SDS, 50 μg/ml proteinase K) at 68°C and 1,300 rpm for 2h. The chromatin was recovered in the supernatant and then DNA was extracted using Phenol:Chloroform:Isoamyl Alcohol (Sigma-Aldrich, ref. P2069). Chromatin was then precipitated with 3 M NaAc (1/10 of total volume), 1 μl Glycogen (stock 20 mg/ml), and 2.5 volumes of pre-chilled EtOH 100% over night at -20°C. The next day, samples were centrifuge at 4°C at 13,000 rpm for 30 min. Pellets were washed once in EtOH 70% and pellets were let to dry before resuspension in DNAse-free water. Of note, inputs were also extracted to serve as reference. Finally, a q- PCR was performed using specific primers for the promoters of IL-1β, IL-6, and IL-10 **(Supplementary table 1)**.

### Western Blot

Macrophages were lysed with 100 μl of lysis buffer (20 mM Tris pH 7.5, 150 mM NaCl, 0.5% NP-40, 50 mM NaF and 1 mM sodium orthovanadate supplemented with complete protease inhibitor cocktail) at the indicated time post HRV16 exposure and/or TLR stimulation, by incubating on ice 5min then vortexing for a total of 3 times. Lysates were centrifuged at 13,000 rpm for 10 min at 4 °C. The supernatants were collected and stored at −20 °C. Protein were quantified by BCA (BCA dosage kit, Pierce ref. 23225). Similar quantity of proteins for each condition was analyzed by SDS-PAGE. Proteins were transferred onto a polyvinylidene difluoride membrane (Millipore, ref. IPVH00010) at 4 °C for 100 min at 100V and incubated in blocking solution (0.1% Tween-20 supplemented with 5% milk or BSA in TBS 1X) for 2h. Blots were rinsed with TBS 0.1% Tween-20, and primary antibody against p65-phosphorylated (1/1000 dilution, Cell signaling, ref. 3033), IRF3-phosphorylated (1/1000 dilution, Cell Signaling, ref. 4947S), SIRT4 (1/1000 dilution, Cell signaling ref. 69786), TIP60 (1/500 dilution, Bio-Techne ref.NBP2-20647), Histone Deacetylase (HDAC) Antibody Sampler Kit (Ozyme ref.9928T), GAPDH (1/1000 Cell Signaling Techonology, ref.97166S) was incubated in the blocking solution overnight. The membrane was further washed and incubated with HRP-coupled secondary antibodies anti-mouse IgG (1/10 000 dilution, Jackson Immunoresearch, ref. 715) or ant-rabbit IgG (1/10 000 dilution, Jackson Immunoresearch, ref. 711-035-152) in blocking buffer for 45 min. Detection was performed using ECL Dura Substrate (GE Healthcare, ref. 34075), bands were imaged with a Fusion (Vilber Lourmat) and quantified in ImageJ.

### Immunofluorescence Microscopy

Cells were fixed in 4% paraformaldehyde for 15 min at room temperature then washed in PBS. Cells were then permeabilized 10min with PBS 1X, 2% FBS, 0.05% saponin (Sigma-Aldrich, S7900) for IL-6 and GM-130 staining or 2min with PBS + 0,1% triton for phosphorylated p65 staining. Coverslips were incubated with primary antibodies: phosphorylated-p65 (1/1000 dilution, Cell signaling Ref. 3033), GM-130 (1/500 dilution, BD ref.610822) or IL-6 (1/200 dilution, Invitrogen ref.701028) diluted in PBS 1X, 2% FBS, with 0.05% saponin (Sigma-Aldrich, S7900) for IL-6 and GM-130 staining, for 45 min at room temperature. Coverslips were washed 3 times then incubated with the following secondary antibodies for 30 min at room temperature: anti-rabbit-AF488 for phosphorylated-p65 and IL-6 (1/500 dilution, Jeckson Immunoresearch ref.711-546-152) or anti-mouse-Cy3 for GM- 130 (1/500 dilution, Jeckson Immunoresearch ref.715-166-151). Nucleus were stained with DAPI (1/30 000 dilution, Sigma ref. D9542). After 3 washes, coverslips were mounted onto slides with Fluoromount-G (INTERCHIM SA, ref. FP-483331).

Images for phosphorylated-p65 staining were acquired using an inverted wide-field microscope (Leica DMI6000) equipped with a 63× (1.4 NA) HCX APO oil-immersion objective, an ORCA flash 4 LT camera (Hamamatsu), and a HXP R 120W/45C VIS lamp. A z-stack series of images was captured at intervals of 0.3 µm. For the analysis, the total number of nuclei was counted, and DAPI/GFP overlays were used to determine the number of p65 (GFP)-positive nuclei. The percentage of p65-positive nuclei was calculated. The mean percentage was determined from 4 to 6 view fields per condition and normalized to the Mock condition.

For IL-6 and GM130 staining, images were acquired with the same inverted microscope (Leica DMI6000) but fitted with a confocal Spinning Disc scanning head (Yokogawa CSU- X1M1). This system included a 63× Plan APO oil-immersion lens (1.4 NA) and a motorized XY stage (Märzhäuser Wetzlar SCAN IM 127-83). Z-stack images were captured at 0.3 µm intervals along the optical axis of the microscope. For the analysis, a Z-projection of IL- 6 images was performed to measure total intensity. The number of nuclei was counted to normalize the staining intensity as a ratio per cell. The average ratio was then calculated from 3 to 4 view fields and compared to the Mock condition. Additionally, channels were merged, and images were converted to RGB. A consistent threshold was applied across all experiments to identify colocalized signals. Both the colocalized area and the total labeled area were measured, and a percentage of colocalized area was calculated. The average percentage was derived from 3 to 4 view fields and compared to the Mock condition. MetaMorph 7.7.5 (Molecular Devices) software was used for system control and image acquisition. All images were analyzed using ImageJ (Version Java 21.0.7, 64-bit).

### FACS staining

Cells were washed 2 times with PBS and detached using cell lifter in 1 mL of FACS buffer (PBS + 2% FBS). Cells were then equally distributed in wells of a 96 well-plate with round bottom (Falcon, ref. 351177). Cells were centrifuged at 1,200 rpm for 5min, supernatant was removed and cells placed on ice and resuspended in 100 μL of FACS buffer. Proper quantities of antibodies were added except for 1 well per condition to serve as the unstained control. The following antibodies were used: IgG2a-PE (20 μl/wells, Beckton Dickison, ref.555574), CD71-PE (20 μl/wells, Beckton Dickison, ref.555537). Cells were incubated on ice for 45min in the dark and then centrifuged at 1,200 rpm 4°C for 5min. Supernatant was removed and cells resuspend in 100 μL of FACS buffer to wash the cells. This step was repeated a second time then cells were fixed for 45min in the dark on ice with 100 μL of 4% paraformaldehyde. Cells were then wash as before twice and finally resuspended in 300μL of FACS buffer. Cells were analyzed using a Attune NxT Flow Cytometer, using the 488-laser acquiring 10.10^4^ events per minutes. Percentage of positive cells was determined using FlowJo software (Version 10.10).

### Graphics and statistical analysis

Graphics and statistical analysis were done using GraphPad Prism (version 11.0.1). Statistical analysis performed for each graph is indicated in the Figures’ legends.

## Results

### HRV16 differentially inhibits IL-6, IL-1β and IL-10 production in macrophages

Previous work from the team demonstrated that HRV16 exposure impairs macrophage responses to secondary bacterial infection^25^. To further investigate the underlying mechanisms, we established a simplified model of bacterial superinfection in which hMDMs were exposed to HRV16 and subsequently stimulated with LPS to activate TLR4 signaling (**Figure 1A**).

**Figure 1:**
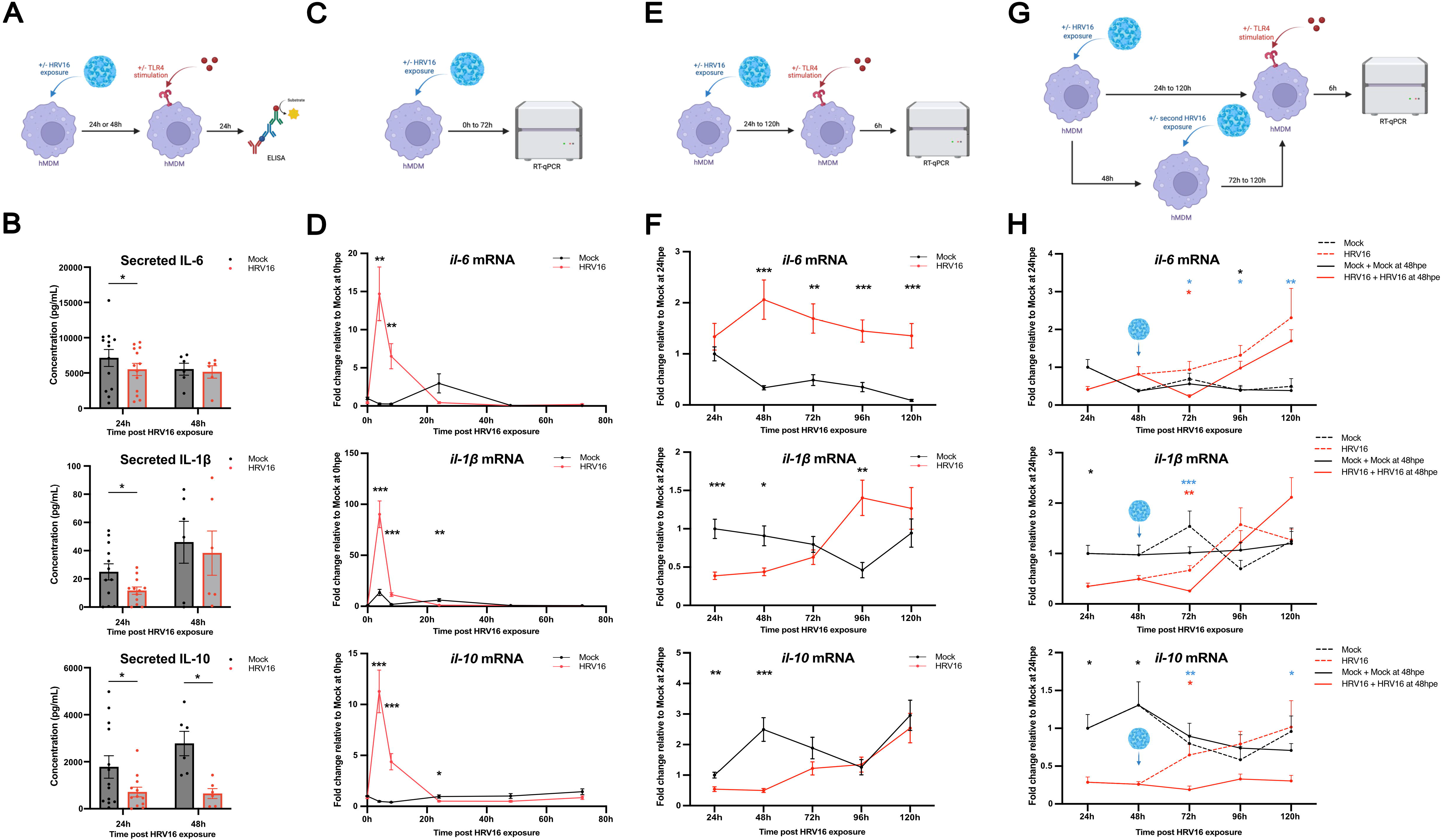
HRV16 differentially inhibits IL-6, IL-1β and IL-10 production in macrophages. **(A-B)** Human monocyte-derived macrophages (hMDMs) were exposed (HRV16) or not (Mock) to 1×10^7^ TCID50/mL of HRV16. At 24h and 48h post exposure (hpe), hMDMs were stimulated with 100 ng/mL of LPS. The supernatants were recovered 24h after stimulation and analyzed by ELISA. **(A)** Experimental protocol. **(B)** IL-6, IL-10 and IL-1β were quantified by ELISA. n=13 at 24hpe and n=6 at 48hpe. Paired T tests were performed. **(C- D)** hMDMs were exposed (HRV16) or not (Mock) to 1×10^7^ TCID50/mL of HRV16. RNAs were extracted at 0hpe, 4hpe, 8hpe, 24hpe, 48hpe, and 72hpe. **(C)** Experimental protocol. **(D)** The mRNA levels for *il-10*, *il-1*β and *il-6* were analyzed by RT-qPCR. n=6, Two-way ANOVA statistical tests were performed. **(E-F)** hMDMs were exposed (HRV16) or not (Mock) to 1×10^7^ TCID50/mL of HRV16. At 24hpe, 48hpe, 72hpe, 96hpe or 120hpe, hMDMs were stimulated with 100 ng/mL of LPS. The RNAs were extracted 6h after stimulation. **(E)** Experimental protocol, **(F)** The mRNA levels for *il-10*, *il-1*β and *il-6* were analyzed by RT-qPCR. n=11, two-way ANOVA statistical tests were performed. **(G-H)** hMDMs were exposed (HRV16) or not (Mock) to 1×10^7^ TCID50/mL of HRV16. At 48hpe, hMDMs were exposed a second time to HRV16 (HRV16 + HRV16 48hpe) or not (Mock + Mock 48hpe). At 24h, 48h, 72h, 96h or 120h post the first exposure, hMDMs were stimulated with 100 ng/mL of LPS and the RNAs were extracted 6h after stimulation. **(G)** Experimental protocol. **(H)** The mRNA levels for *il-10*, *il-1*β and *il-6* were analyzed by RT- qPCR. n=6. Two-way ANOVA statistical tests were performed. Black stars: significative difference between « HRV16 » and « Mock »; red stars: significative difference between « HRV16 + HRV16 at 48 hpe » and « HRV16 »; blue stars: significative difference between « HRV16 + HRV16 at 48 hpe » and « Mock + Mock at 48 hpe ». All data are represented as the mean +/- the standard error of the mean (SEM). Statistical significance is indicated as follow: *p<0,05; **p<0,01; ***p<0,001.

To confirm cytokine inhibition in this model, hMDMs were exposed to HRV16 and stimulated with LPS either 24h or 48h post viral exposure (hpe). Cytokine secretion was analyzed 24h after stimulation (**Figure 1A**). HRV16-exposed macrophages secreted significantly lower levels of IL-6 (1.3-fold decrease), IL-1β (2.2-fold decrease), and IL-10 (2.5-fold decrease) compared with Mock-treated cells at 24hpe (**Figure 1B**). In contrast to IL-6 and IL-1β, IL-10 secretion remained significantly inhibited at 48hpe with a 4.3-fold decrease. These observations suggested that HRV16 differentially regulates cytokine secretion and that distinct mechanisms may account for the transient inhibition of IL-6 and IL-1β compared with the more sustained inhibition of IL-10.

To determine whether cytokine inhibition occurred at the transcriptional level, we first analyzed these cytokines mRNA expression at multiple time points following HRV16 exposure in the absence of LPS stimulation (**Figure 1C**). All 3 cytokines displayed an early induction peak at 4hpe that rapidly declined. By 24hpe, IL-10 and IL-1β mRNA levels were significantly reduced compared with Mock-treated cells, while IL-6 expression showed a similar downward trend (**Figure 1D**). To assess if similar results would be observed upon bacteria exposure, we next investigated cytokine transcription following LPS stimulation at the same times post HRV16 exposure (**Figure 1E**). Interestingly, IL-6 mRNA levels were increased approximately 2-fold in HRV16-exposed macrophages at most time points (**Figure 1F**). In contrast, IL-10 and IL-1β transcriptions were significantly reduced at 24hpe and 48hpe, but this inhibition progressively disappeared at later time points. Notably, IL-1β expression was even increased at 72hpe compared with Mock-treated cells. Together, these data indicate that HRV16 differentially regulates cytokine expression. While IL-10 and IL-1β undergo transient transcriptional inhibition, IL-6 secretion is impaired despite preserved or increased transcriptional induction, suggesting the involvement of a post-transcriptional mechanism.

In our model, hMDMs are exposed to HRV16 only during the initial infection, since we previously showed that HRV16 does not replicate in macrophages^24^. However, in patients, HRV16 will be continuously produced by permissive cells (e.g. lung epithelial cells)^28,29^. Therefore, *in vivo*, macrophages are likely repeatedly exposed to newly produced viral particles during the infection. To better mimic this physiological context, macrophages were exposed to a second dose of HRV16 at 48hpe before LPS stimulation (**Figure 1G**). Interestingly, following this second exposure, IL-6 transcription was significantly reduced at 72hpe but rapidly recovered at later time points (**Figure 1H**). IL-1β transcription also displayed prolonged but transient inhibition after the second exposure. In contrast, IL-10 transcriptional inhibition was maintained up to 120hpe with a sustained 2- to 4-fold decrease (**Figure 1H**) suggesting that additional transcriptional repressive mechanisms may arise upon repeated viral exposure.

Altogether, these results demonstrate that HRV16 inhibits macrophage cytokine production through distinct mechanisms depending on the cytokine studied. IL-10 and IL- 1β undergo transcriptional repression, whereas IL-6 secretion appears to be primarily regulated at the post-transcriptional level.

### HRV16 promotes intracellular retention of IL-6 by disrupting intracellular trafficking

Given that IL-6 secretion was impaired despite increased transcriptional induction, we next investigated whether HRV16 interfered with intracellular trafficking and protein secretion pathways. Previous work from the laboratory demonstrated that HRV16 disrupts Golgi apparatus organization in macrophages^23^. We therefore hypothesized that HRV16 may impair cellular trafficking including Golgi apparatus-dependent protein secretion pathways and consequently inhibit cytokine secretion.

By immunofluorescence, we investigated whether IL-6 accumulated intracellularly following HRV16 exposure. Indeed, we observed significantly increased levels of stained intracellular IL-6 (1.3-fold increase) at 48hpe in HRV16-exposed macrophages compared to Mock (**Figure 2A and 2B**), confirming cellular retention. Interestingly, we observed that IL-6 seemed to accumulate in clusters near the Golgi apparatus (**Figure 2A, cropped cell 4 and Figure 2B**).

**Figure 2:**
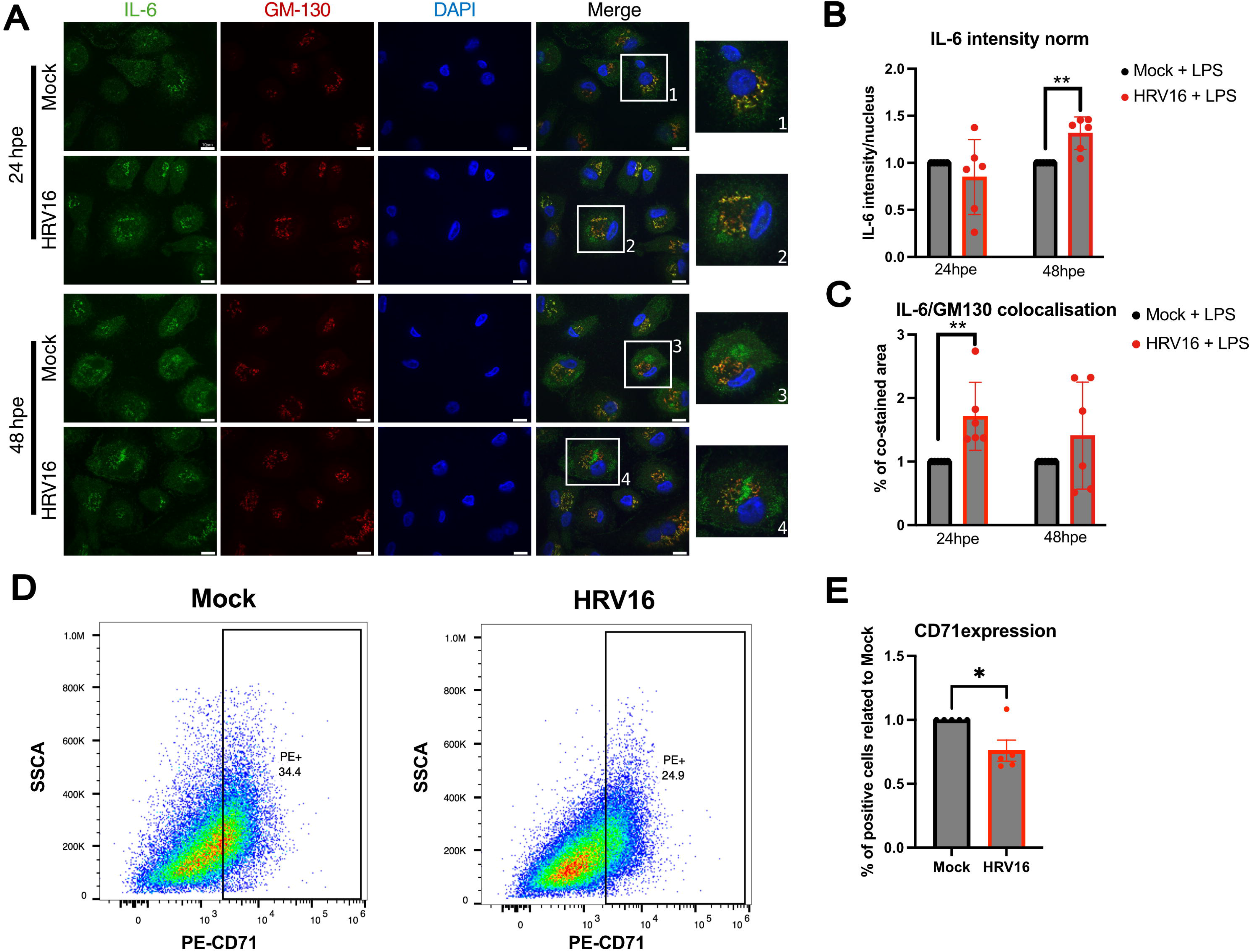
HRV16 promotes intracellular retention of IL-6 by disrupting intracellular trafficking. **(A-C)** hMDMs were exposed (HRV16) or not (Mock) to 1×10^7^ TCID50/mL of HRV16. At 24hpe or 48hpe, hMDMs were stimulated with 100 ng/mL of LPS for 24h. After stimulation, cells were fixed and stained by immunofluorescence with an antibody against IL-6 and GM-130 (cis Golgi Apparatus) as well as DAPI. **(A)** Representative images of the staining. **(B)** Intracellular IL-6 levels and **(C)** IL-6/Golgi apparatus colocalization were quantified using ImageJ. n=6. **(D-E)** hMDMs were exposed (HRV16) or not (Mock) to 1×10^7^ TCID50/mL of HRV16. At 24hpe cells were stained and fixed for flow cytometry analysis with an antibody against CD71. **(D)** Representative FACS plot showing the percentage of positive cells for CD71 membrane expression. (**E)** Percentage of positive cells for CD71 staining related to Mock condition. **(B-C, E)** T-test statistical test was performed. Data are presented as the mean +/- the standard error of the mean (SEM). Statistical significance is indicated as follow: *p<0,05; **p<0,01.

To determine whether IL-6 was retained in the Golgi apparatus, we analyzed the colocalization between IL-6 and GM-130, a marker of the cis-Golgi apparatus. HRV16- exposed macrophages displayed 1.7-fold increased IL-6 colocalization with the Golgi apparatus at 24hpe (**Figure 2A and 2C**). These results suggest that IL-6 remains trapped inside the Golgi apparatus at early time points following viral exposure and is then retained in the cytoplasm.

To evaluate if HRV16 only impacted Golgi apparatus associated secretion or further steps of internalization and recycling, we analyzed the membrane expression of CD71, a known marker for intracellular trafficking as it is repeatedly internalized and recycled through endosomes^30^. HRV16-exposed macrophages displayed a 1.3-fold decreased plasma membrane expression of CD71 compared with Mock-treated cells (**Figure 2D and 2E**). This result indicates a defect in intracellular trafficking and exocytosis, as also suggested by our previous results ^23^.

Together, these results indicate that HRV16 induces intracellular retention of IL-6 and impaired cytokine secretion in macrophages.

HRV16 induces global epigenetic remodeling in macrophages

Subsequently, to investigate the mechanisms underlying cytokine transcriptional inhibition, we examined whether HRV16 exposure induced epigenetic alterations in macrophages.

We first analyzed global DNA methylation levels on two classical methylation sites: 5mC and 5hmC. No significant differences were found in 5mC nor 5hmC levels between HRV16-exposed and Mock-treated macrophages, either in basal conditions or following LPS stimulation (**Figure 3A**).

**Figure 3:**
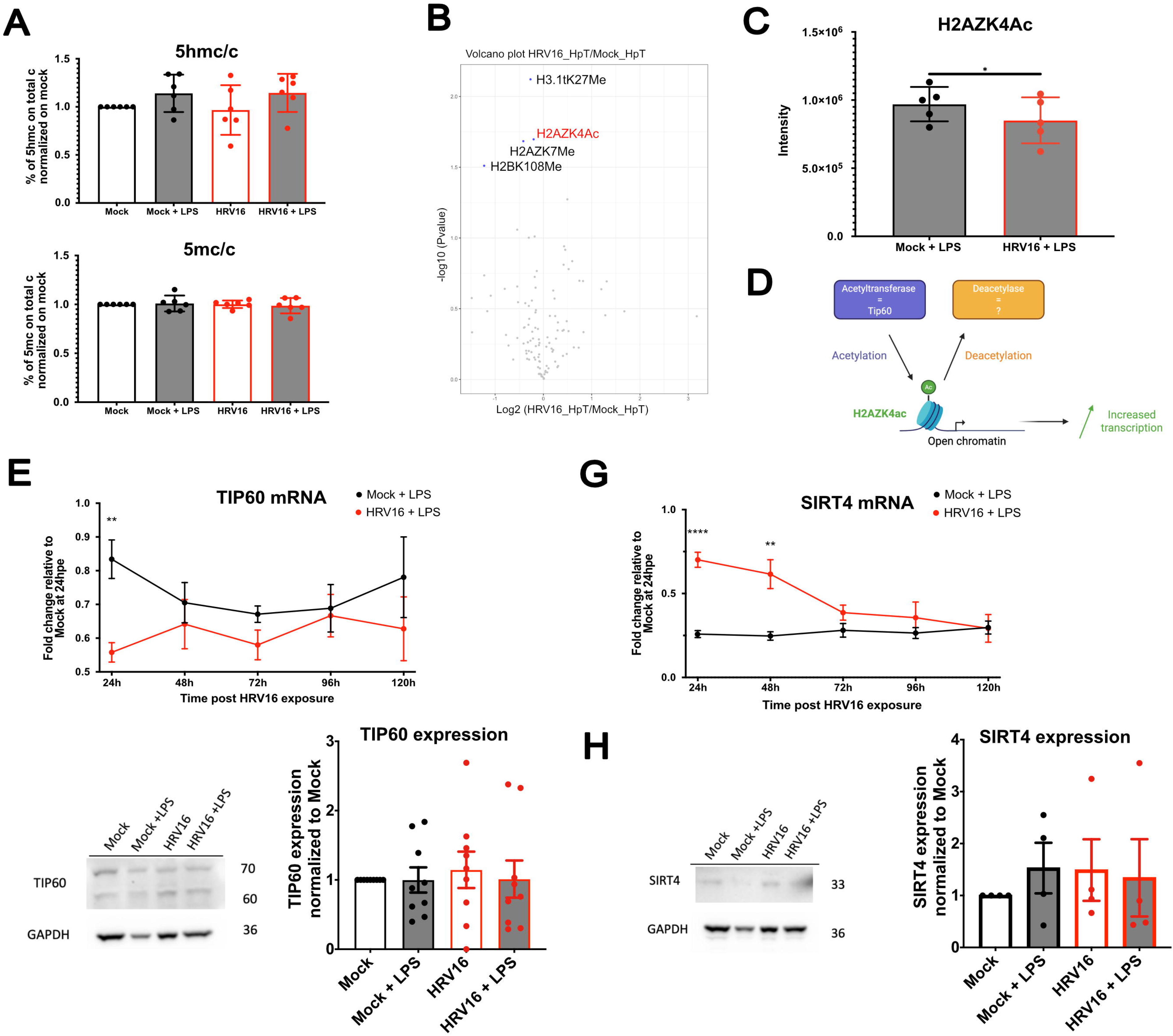
HRV16 induces epigenetic remodeling in macrophages. **(A-C, F-H)** hMDMs were exposed (HRV16) or not (Mock) to 1×10^7^ TCID50/mL of HRV16. At 24hpe hMDMs were stimulated or not with 100 ng/mL of LPS for 24h. **(A)** DNA was extracted and digested. Levels of 5hmc/c and 5mc/c were then analyzed by mass spectrometry. n=6. One-way ANOVA statistical tests were performed. **(B)** Histones modifications were analyzed by mass spectrometry. n=5. Intensities of HRV16 conditions were compared to Mock ones by a welch test. P-value threshold was set to 0.05. **(C)** H2AZK4Ac expression from the mass spectrometry analysis. n=5. Paired T-test statistical analysis was performed. **(D)** Schematic representation of H2AZK4Ac regulation. **(E, G)** hMDMs were exposed (HRV16) or not (Mock) to 1×10^7^ TCID50/mL of HRV16. At 24hpe, 48hpe, 72hpe, 96hpe or 120hpe, hMDMs were stimulated with 100 ng/mL of LPS. The RNAs were extracted 6h after stimulation. The mRNA levels for **(E)** *tip60* and **(G)** *sirt4* were analyzed by RT-qPCR. n=6, Two-way ANOVA statistical tests were performed. **(F, H)** The proteins were extracted 24h after stimulation and the expression of **(F)** TIP60 (n=9) and **(H)** SIRT4 (n=4) were analyzed by western-Blot. Two-way ANOVA statistical tests were performed. **(A, C)** Data are presented as the mean +/- the standard deviation (SD). **(E-H)** Data are presented as the mean +/- the standard error of the mean (SEM). Statistical significance is indicated as follow: *p<0,05; **p<0,01; ***p<0,001; ***p<0,0001.

We next investigated histone modifications induced by HRV16 using mass spectrometry and identified 11 histone marks whose levels were significantly decreased following HRV16 exposure (**Figure 3B** and **Supplementary table 2**). Among these candidates, H2AZK4Ac is associated with gene transcription activation, and therefore its decrease could contribute to the observed transcriptional inhibition^31^. H2AZK4Ac was decreased by 1.2-fold in HRV16-exposed macrophages (**Figure 3C**).

To determine whether this decrease could result from altered expression of histone-modifying enzymes, we analyzed the expression of TIP60, the histone acetyltransferase involved in H2AZ acetylation^32^, together with several histone deacetylases, as the one involved in H2AZK4 deacetylation as not been identified yet (**Figure 3D**). A 2-fold decrease of TIP60 mRNA expression was observed at 24hpe (**Figure 3E**). However, no significant difference was detected at the protein level (**Figure 3F**). Similarly, several histone deacetylases displayed altered transcriptional profiles (**Figure 3G** and **Supplementary Figure 1A-I**), including SIRT4, which showed 3-fold increased mRNA expression following HRV16 exposure (**Figure 3G**). Yet, no corresponding changes were observed at the protein level (**Figure 3H** and **Supplementary Figure 1J-N**).

Together, these results indicate that HRV16 induces epigenetic remodeling in macrophages. However, these changes do not appear to result from major alterations in the expression levels of the histone-modifying enzymes analyzed.

### HRV16-induced epigenetic changes are not directly associated with cytokine transcriptional inhibition

To determine whether the identified epigenetic modification induced by HRV16 contributed to cytokine repression, we analyzed the presence of H2AZK4Ac at the promoters of IL-6, IL-10 and IL-1β by chromatin immunoprecipitation (**Figure 4A**). We additionally examined two histone marks previously associated with HRV16-mediated regulation of genes in macrophages, namely H3K27Me3 and H3K27Ac^24^ (**Figure 4B-C**).

**Figure 4:**
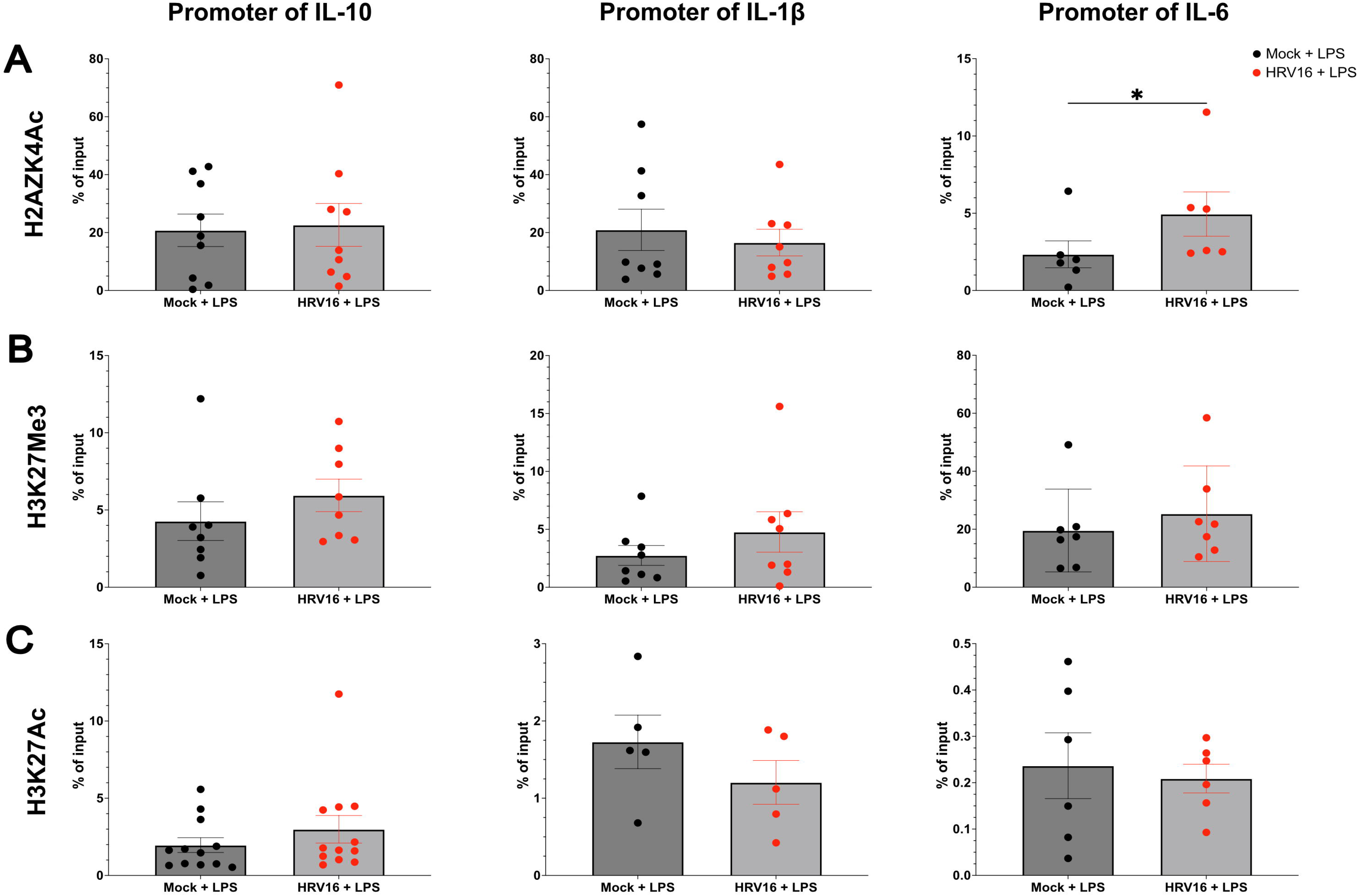
HRV16-induced epigenetic changes are not directly associated with cytokine transcriptional inhibition. **(A-C)** hMDMs were exposed (HRV16) or not (Mock) to 1×10^7^ TCID50/mL of HRV16. At 24hpe, hMDMs were stimulated with 100 ng/mL of LPS for 24h. Then chromatin/proteins were cross-linked and **(A)**H2AZK4Ac, **(B)** H3K27Me3 and **(C)** H3K27Me3 marks were immunoprecipitated. IgG was used as a negative control. The promoters of *il-10*, *il-1*β and *il-6* were analyzed by qPCR. Results are shown as percentage of input and t-tests were performed. Data are presented as the mean +/- the standard error of the mean (SEM). Statistical significance is indicated as follow: *p<0,05.

H2AZK4Ac was not decreased at the promoters of IL-10 or IL-1β following HRV16 exposure. In contrast, the IL-6 promoter displayed increased H2AZK4Ac levels in HRV16- exposed macrophages (**Figure 4A**). Similarly, no significant differences in H3K27Me3 or H3K27Ac levels were detected at the promoters of IL-6, IL-10 or IL-1β after HRV16 exposure (**Figure 4B-C**).

Overall, these results indicate that although HRV16 induces global epigenetic remodeling in macrophages, the histone modifications investigated here do not directly explain the transcriptional repression of IL-10 and IL-1β. In contrast, the upregulation of IL-6 expression correlates with the increased H2AZK4Ac at its promoter, suggesting a selective epigenetic regulation mechanism.

### HRV16 disrupts TLR signaling pathways by inhibiting phosphorylated-p65 translocation to the nucleus

As the epigenetic analyses performed did not identify the cause of cytokine transcriptional repression, we next investigated whether HRV16 altered the upstream TLR signaling pathways.

TLR4 signaling activates both MyD88-dependent and TRIF-dependent pathways leading to the activation of respectively NF-κB and IRF3^9^. To determine whether HRV16-mediated inhibition was specific to TLR4 signaling, macrophages were stimulated with Poly(I:C) to induce TLR3, which mainly activates IRF3 but can also partially signal through NF-κB^12^ (**Figure 5A**). Following TLR3 stimulation, HRV16-exposed macrophages displayed a 2- fold increase of IL-6 mRNA levels, similarly to TLR4-stimulated cells (**Figure 5B**). Whereas IL-1β mRNA levels remained unchanged, IL-10 expression was significantly reduced by 2- fold at 24hpe and 48hpe (**Figure 5B**). Although the results for IL-1β transcription are different between the two stimulations, the inhibition of IL-10 transcription observed with both TLR3 and TLR4 signaling could suggest that HRV16 interferes with either IRF3 or NF-κB dependent transcription activation.

**Figure 5:**
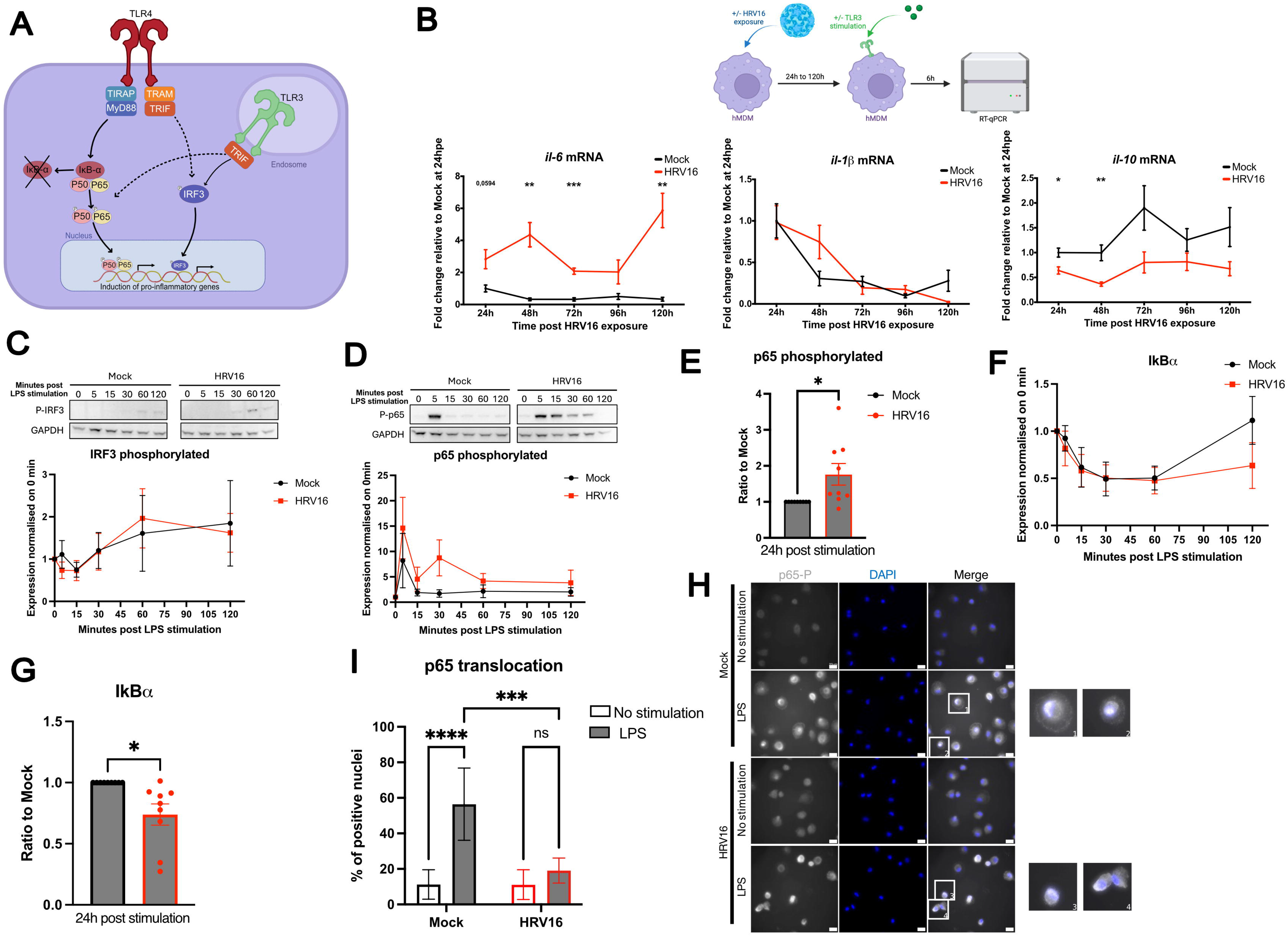
HRV16 disrupts TLR signaling pathways by inhibiting phosphorylated-p65 translocation to the nucleus. **(A)** Schematic representation of TLR4, and TLR3 signaling pathways. **(B)** hMDMs were exposed (HRV16) or not (Mock) to 1×10^7^ TCID50/mL of HRV16. At 24hpe, 48hpe, 72hpe, 96hpe or 120hpe, hMDMs were stimulated with 5 μg/mL of Poly(I:C)-HMW. The RNAs were extracted 6h after stimulation. The mRNA levels for *il-6*, *il-10* and *il-1*β were analyzed by RT-qPCR. n=6, two-way ANOVA statistical tests were performed. **(C-G)** hMDMs were exposed (HRV16) or not (Mock) to 1×10^7^ TCID50/mL of HRV16. At 24hpe, hMDMs were stimulated with 100 ng/mL of LPS **(C,D,F)** for 0min, 5min, 15min, 30min, 1h or 2h or **(E,G)** for 24h. After stimulation, proteins were extracted and the expression of **(C)** phosphorylated-IRF3, **(D-E)** phosphorylated-p65, or **(F-G)** IκBα were analyzed by Western-Blot. **(H-I)** hMDMs were exposed (HRV16) or not (Mock) to 1×10^7^ TCID50/mL of HRV16. At 24hpe, hMDMs were stimulated with 100 ng/mL of LPS for 30min. After stimulation, cells were fixed, permeabilized and stained by Immunofluorescence with an antibody against phosphorylated-p65, as well as DAPI. **(H)** Representative images. **(I)** Phosphorylated p65 translocation was quantified with ImageJ and presented as percentage of positive nucleus. n=5, two-way ANOVA statistical tests were performed. All data are presented as the mean +/- the standard error of the mean (SEM). Statistical significance is indicated as follow: *p<0,05; **p<0,01; ***p<0,001; ***p<0,0001.

Therefore, we next analyzed the activated/phosphorylated status of the transcription factors IRF3 and p65/RelA. Phosphorylated IRF3 levels remained unchanged between Mock and HRV16-exposed macrophages between 0 and 120min post LPS stimulation (**Figure 5C**), suggesting that IRF3 signaling was preserved. However, an increase of the phosphorylated p65 was observed in HRV16-exposed macrophages at early (0 to 120 min post stimulation) and late (24h post stimulation), with a significant 2-fold increase at 24h post stimulation (**Figure 5D-E**). As NF-κB negative feedback loop and return to homeostasis depend on the resynthesis of its cytoplasmic inhibitor, IκB-α, which normally binds and sequesters NF-κB dimers to prevent their nuclear translocation^11^, we assessed IκB-α expression after HRV16 exposure. A reduced IκB-α expression was visible at 120min and 24h post LPS stimulation in the HRV16 condition compared to the Mock (**Figure 5F-G**), suggesting a dysregulation of NF-κB activity.

To confirm that HRV16 could block p65 activity, we quantified the nuclear translocation of phosphorylated p65. As expected, we observed more phosphorylated p65 in the LPS treated cells compared to the non-stimulated (**Figure 5H**). Interestingly, we could observe perinuclear accumulations of phosphorylated-p65 in the HRV16 condition (**Figure 5H, cropped cell 4**). HRV16-exposed macrophages displayed a 3-fold reduction in number of phosphorylated p65 positive nuclei compared with Mock-treated cells (**Figure 5H-I**).

Together, these results demonstrate that HRV16 impairs NF-κB p65/RelA nuclear translocation while preserving IRF3 signaling. This mechanism likely contributes to the selective inhibition of NF-κB-dependent cytokine transcription observed following TLR4 and TLR3 stimulation.

## Discussion

In this study, we investigated the mechanisms by which HRV16 impairs macrophage cytokine secretion. We focused on HRV16, a HRV-A serotype, that is most frequently associated with COPD exacerbations^18,19^. Our findings reveal that HRV16 differentially regulated macrophage cytokine secretion through distinct transcriptional and post-transcriptional mechanisms. Whereas IL-10 and IL-1β were inhibited at both the transcriptional and secretion levels, IL-6 transcription was increased despite reduced cytokine secretion (**Figure 6**). Together, these findings supported a model in which HRV16 modulates macrophage cytokine response through multiple mechanisms.

**Figure 6:**
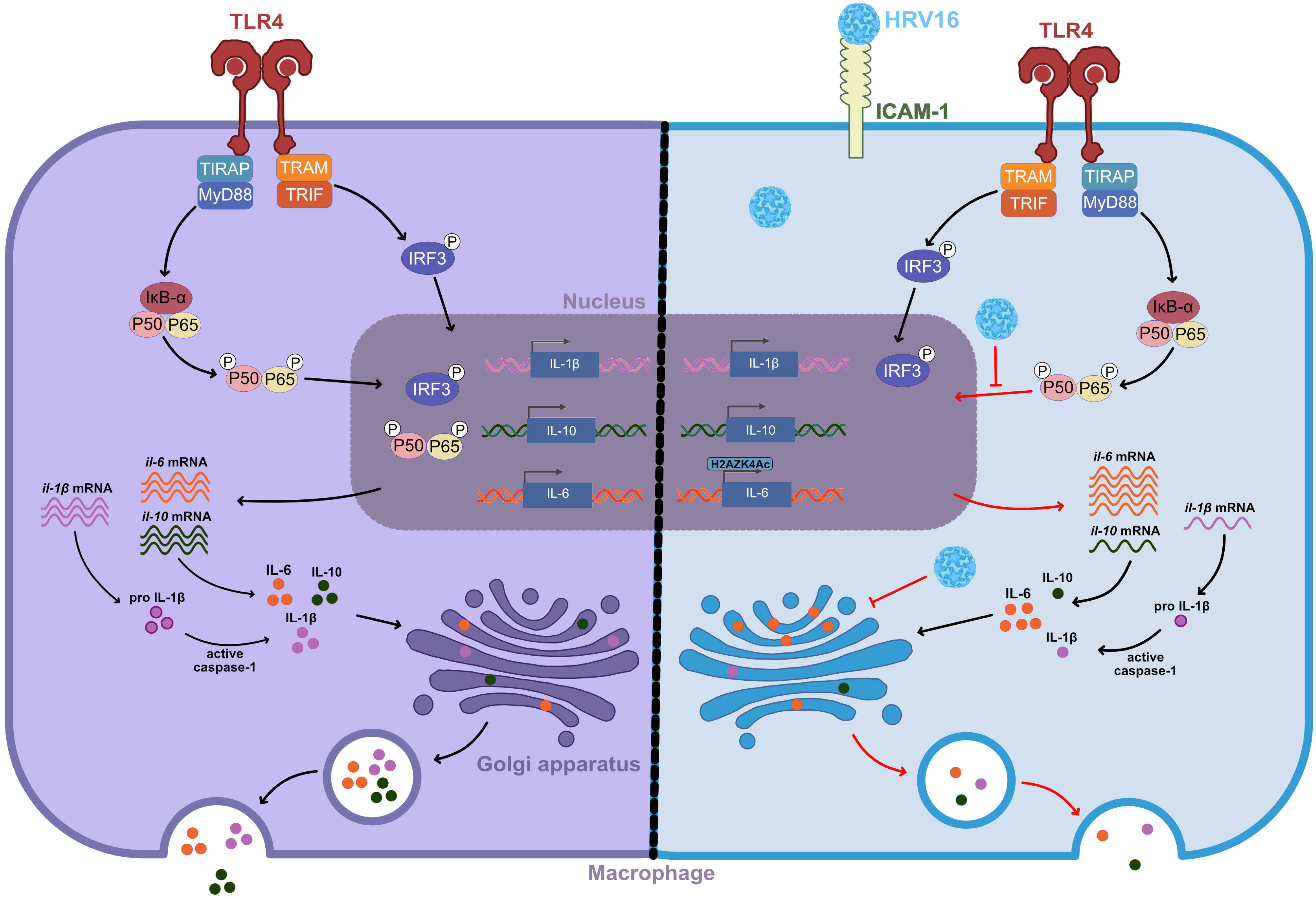
Graphical abstract. Left side: Normal macrophages response to a stimulation with lipopolysaccharide (LPS). LPS binds to TLR4 and activates both the MyD88-dependent and TRIF-dependent signaling pathways. The MyD88 pathway promotes IκBα degradation, leading to NF-κB (p65/p50 dimer) phosphorylation and nuclear translocation, whereas the TRIF pathway induces IRF3 phosphorylation and nuclear translocation. Once in the nucleus, these transcription factors drive the expression of inflammatory genes including IL-1β, IL-10, and IL-6. The resulting mRNAs are translated into proteins in the cytoplasm, except for IL-1β which will be first translated to a pro form before being cleaved by the activated caspase-1. Cytokines are subsequently processed through the Golgi apparatus and secreted via the conventional exocytosis pathway. Right side: Macrophages responses after an exposition to HRV16. HRV16 inhibits the nuclear translocation of phosphorylated p65, resulting in reduced transcription of IL-1β and IL-10. In contrast, IL-6 transcription is enhanced due to the HRV16-dependent enrichment of the activating histone mark H2AZK4Ac at the promoter of IL-6. In addition, HRV16 induces IL-6 retention within the cis-Golgi, impairing cytokine trafficking and secretion. Black arrow: normal functioning process; Inhibitory arrows: step inhibited by the virus; Red arrows: downstream consequences of the virus inhibition.

The increase of IL-6 transcription levels despite reduced cytokine secretion suggested the existence of a post-transcriptional mechanism. The observed intracellular accumulation of IL-6 together with reduction of CD71 membrane expression indicated that HRV16 impairs intracellular protein trafficking. These results are also in line with previous work from our group that showed a defect in intracellular trafficking affecting phagosome maturation of macrophages, due to ARL5b upregulation induced by HRV16 in macrophages, where fluorescence and electron microscopy revealed perturbation of the Golgi and endosomal compartments^23^. However, another study also investigating the effects of HRV16 on the Golgi apparatus fragmentation reported that protein secretion was not affected by said disruption^33^. Unlike our study, these experiments were performed in HeLa cells, which support productive viral infection, whereas macrophages do not. The viral proteins identified as responsible for Golgi apparatus fragmentation were the non-structural proteins HRV16 3A, 3AB, and 2B. In our model, due to the absence of productive replication in human monocyte-derived macrophages, the structural proteins delivered with the viral particles are likely to be responsible for the observed effects. Unfortunately, we were unable to directly assess their contribution due to technical limitations associated with primary macrophages. Nevertheless, it is important to note that HRV16 possesses a positive-sense RNA genome, meaning that viral entry into macrophages could result in immediate translation of the entire viral polyprotein, including both structural and non-structural proteins, even in the absence of productive virion formation. Under such conditions, macrophages could transiently express HRV16 3A, 3AB, and 2B proteins, potentially contributing to the Golgi apparatus disruption observed here.

Our study also does not explore the possibility that HRV16 affects other stages of the secretory pathway, including protein translation, endoplasmic reticulum export, or vesicle maturation. Future studies will therefore be necessary to identify the precise mechanisms responsible for HRV16-induced cytoplasmic cytokine retention. Nevertheless, our observations clearly demonstrate that HRV16 targets macrophage secretory pathways.

Furthermore, replication of other Picornaviridae family members, including poliovirus^34,35^ and coxsackievirus^36^ has been shown to inhibit protein secretion through disruption of intracellular trafficking. However, these studies were also made in cell types permissive to viral replication. Together, our and other findings suggest that disruption of intracellular trafficking may represent a broader feature of HRV16 and, more generally, of Picornaviridae infections. In macrophages, this phenomenon appears to contribute directly to the inhibition of cytokine secretion.

We next investigated the upstream signaling pathways regulating cytokine transcription. Our data identified impaired NF-κB p65 nuclear translocation as the central mechanism underlying HRV16-mediated cytokine repression. Interestingly, HRV16 did not prevent p65 phosphorylation but specifically reduced its nuclear translocation, suggesting that the virus uncouples NF-κB activation from its translocation. Such a dissociation could result from disruption of the cytoskeleton-dependent trafficking pathways required for NF-κB translocation^37^. This hypothesis is consistent with our observations of IL-6 cytoplasmic retention resulting from impaired intracellular trafficking. However, the precise molecular mechanism remains to be determined.

The pathway-specific effects observed following TLR stimulation further support the role of NF-κB nuclear translocation defect in the transcriptional inhibition of Il-10 and Il-1β. Although TLR3 signaling activates both NF-κB and IRF3, IRF3 represents its dominant transcriptional effector^12^. Consistent with this, HRV16 did not alter IRF3 activation, suggesting that preserved IRF3 signaling may partially compensate for defective NF-κB activity and thereby explain the more limited transcriptional inhibition observed following TLR3 stimulation.

Interestingly, while IL-10 and IL-1β transcription was inhibited, IL-6 transcription remained preserved, suggesting that IL-10 and IL-1β are more sensitive to impaired NF-κB nuclear translocation than IL-6. One possible explanation is the increased enrichment of the activating epigenetic mark H2AZK4Ac at the IL-6 promoter. Such enrichment could maintain a more accessible chromatin configuration, allowing the fraction of NF-κB that still reaches the nucleus to preferentially interact with the IL-6 promoter and sustain its transcription. These findings therefore suggest a previously unrecognized role for H2AZK4Ac as an immune gene-associated epigenetic mark.

Among the histone modifications identified in our global proteomic analysis, we focused on H2AZK4Ac as it is an active epigenetic mark and an antibody was available in human to study this mark. In contrast, the other significantly decreased histone modifications identified were repressive methylation marks, whose decrease would not be able to explain the decreased transcription of IL-10 and IL-1β. However, several limitations of our epigenetic analyses should be acknowledged. Primary human macrophages exhibit substantial inter-donor variability. In our study, we considered marks to be significatively different if they were detected in least 4 out of 5 donors. Due to the variability inherent to our model, it is possible that we overlooked some marks that were detected in 3 or less donors. Moreover, only a fraction of cells is likely infected following HRV16 exposure. This proportion cannot currently be quantified because HRV16 does not replicate in macrophages and no suitable immunofluorescence tools are available to identify infected cells. Consequently, infection-induced chromatin remodeling occurring specifically within infected cells may have been diluted in bulk population analyses. In addition, the only commercially available ChIP-grade antibody recognizing the human H2AZK4Ac mark also recognizes H2AZK7Ac and H2AZK11Ac. Since our proteomic analysis did not detect significant changes in H2AZK7Ac or H2AZK11Ac, subtle alterations specifically affecting H2AZK4Ac may therefore have been underestimated during our ChIP analysis.

Finally, we decided to investigate H3K27Ac and H3K27Me3 even though they were not identified as significantly modified in our proteomic analysis, because previous work reported HRV-induced remodeling of H3K27 marks at promoters of HRV16-regulated genes^24^. However, promoter-specific ChIP analyses did not support a direct association between either modifications and the transcriptional regulation of IL-10, IL-1β, or IL-6, suggesting that these marks are unlikely to account for the selective transcriptional effects observed in our model. However, these results highlight the fact the HRV16 does induce epigenetic modifications in macrophages, and even though these modifications were not directly linked with cytokine secretion inhibition, this shows a more global effect of HRV16 on macrophages’ epigenome and potentially on their functions.

Altogether, our findings demonstrate that HRV16 differentially regulates macrophage cytokine production through both transcriptional and post-transcriptional mechanisms. While defects in intracellular trafficking contribute to impaired cytokine secretion, defective NF-κB p65 nuclear translocation is associated with selective immune genes transcriptional repression. These findings provide new insight into the molecular basis of defective innate immune responses during rhinovirus infection.

---

**Supplementary table 1: List of primers sequences used for q-PCR**

**Supplementary table 2: Proteomic analysis of histone epigenetic modifications induced by HRV16 exposure**

**Supplementary figure 1: Expression of histone deacetylases after HRV16 exposure** (**A-I**) hMDMs were exposed (HRV16) or not (Mock) to 1×10^7^ TCID50/mL of HRV16. At 24hpe, 48hpe, 72hpe, 96hpe or 120hpe, hMDMs were stimulated with 100 ng/mL of LPS. The RNAs were extracted 6h after stimulation. The mRNA levels for **(A)** *sirt1***, (B)** *sirt3***, (C)** *sirt5***, (D)** *sirt7***, (E)** *hdac1*, (**F**) *hdac2*, (**G**) *hdac3*, (**H**) *hdac4*, and **(I)** *hdac6* were analyzed by RT-qPCR. n=6, Two-way ANOVA statistical tests were performed. **(J, N)** The proteins were extracted 24h after stimulation and the expression of **(J)** HDAC1, (**K**) HDAC2, (**L**) HDAC3, (**M**) HDAC4, and **(N)** HDAC6 (n=9) were analyzed by western-blot. Two-way ANOVA statistical tests were performed. Data are presented as the mean +/- the standard error of the mean (SEM). Statistical significance is indicated as follow: *p<0,05; **p<0,01; ***p<0,001.

## Acknowledgements

The authors thank all our funders. The authors thank Chiara Pompili for her help with flow cytometry analysis. Experimental schemes were made with Biorender.

## Authors contribution

Conceptualization: F.N., S.F-D.

Data curation: Z.F-D., Z.M., L.P., M.L., F.B., P.B.A., S.F-D.

Formal analysis: Z.F-D., Z.M., L.P., M.L., F.B., P.B.A., S.F-D.

Funding acquisition: Z.F-D., F.N., S.F-D. Visualization: Z.F-D., M.L., S.F-D. Writing- original draft: Z.F-D., S.F-D.

## Data availability statement

The data that support the findings of this study are available from the corresponding author, S.F-D., upon request.

## References

1. Guan, F. et al. Tissue macrophages: origin, heterogenity, biological functions, diseases and therapeutic targets. Sig Transduct Target Ther 10, 93 (2025).

2. Akira, S., Uematsu, S. & Takeuchi, O. Pathogen recognition and innate immunity. Cell 124, 783–801 (2006).

3. Depierre, M., Jacquelin, L. & Niedergang, F. Phagocytosis. in Encyclopedia of Cell Biology (Second Edition) (eds Bradshaw, R. A., Hart, G. W. & Stahl, P. D.) 286–295 (Academic Press, Oxford, 2023). doi:10.1016/B978-0-12-821618-7.00038-9.

4. Muntjewerff, E. M., Meesters, L. D. & van den Bogaart, G. Antigen Cross-Presentation by Macrophages. Front. Immunol. 11, (2020).

5. Malainou, C., Abdin, S. M., Lachmann, N., Matt, U. & Herold, S. Alveolar macrophages in tissue homeostasis, inflammation, and infection: evolving concepts of therapeutic targeting. J Clin Invest 133, (2023).

6. Pervizaj-Oruqaj, L., Ferrero, M. R., Matt, U. & Herold, S. The guardians of pulmonary harmony: alveolar macrophages orchestrating the symphony of lung inflammation and tissue homeostasis. European Respiratory Review 33, (2024).

7. Duan, T., Du, Y., Xing, C., Wang, H. Y. & Wang, R.-F. Toll-Like Receptor Signaling and Its Role in Cell-Mediated Immunity. Front Immunol 13, 812774 (2022).

8. Fernandes-Santos, C. & de Azeredo, E. L. Innate Immune Response to Dengue Virus: Toll-like Receptors and Antiviral Response. Viruses 14, 992 (2022).

9. Cheng, Z., Taylor, B., Ourthiague, D. R. & Hoffmann, A. Distinct single-cell signaling characteristics are conferred by the MyD88 and TRIF pathways during TLR4 activation. Sci Signal 8, ra69 (2015).

10. Chen, Y., Lin, J., Zhao, Y., Ma, X. & Yi, H. Toll-like receptor 3 (TLR3) regulation mechanisms and roles in antiviral innate immune responses. J Zhejiang Univ Sci B 22, 609–632 (2021).

11. Kawai, T. & Akira, S. Signaling to NF-κB by Toll-like receptors. Trends in Molecular Medicine 13, 460–469 (2007).

12. Chen, Y., Lin, J., Zhao, Y., Ma, X. & Yi, H. Toll-like receptor 3 (TLR3) regulation mechanisms and roles in antiviral innate immune responses. J Zhejiang Univ Sci B 22, 609–632 (2021).

13. Locatelli, M. & Faure-Dupuy, S. Virus hijacking of host epigenetic machinery to impair immune response. J Virol 97, e00658–23.

14. Chen, S., Yang, J., Wei, Y. & Wei, X. Epigenetic regulation of macrophages: from homeostasis maintenance to host defense. Cell Mol Immunol 17, 36–49 (2020).

15. Fremont-Debaene, Z. & Faure-Dupuy, S. Macrophage makeover extreme viral edition: mechanisms of immune subversion and therapeutic perspectives. J Gen Virol 107, 002228 (2026).

16. Faure-Dupuy, S., Durantel, D. & Lucifora, J. Liver macrophages: Friend or foe during hepatitis B infection? Liver Int 38, 1718–1729 (2018).

17. Human Rhinovirus A16 - an overview | ScienceDirect Topics. https://www.sciencedirect.com/topics/medicine-and-dentistry/human-rhinovirus-a16.

18. George, S. N. et al. Human rhinovirus infection during naturally occurring COPD exacerbations. European Respiratory Journal 44, 87–96 (2014).

19. Matsumoto, K. & Inoue, H. Viral infections in asthma and COPD. Respiratory Investigation 52, 92–100 (2014).

20. Sim, Y. S. et al. COPD Exacerbation-Related Pathogens and Previous COPD Treatment. Journal of Clinical Medicine 12, 111 (2022).

21. Viniol, C. & Vogelmeier, C. F. Exacerbations of COPD. Eur Respir Rev 27, 170103 (2018).

22. Jubrail, J. et al. Arpin is critical for phagocytosis in macrophages and is targeted by human rhinovirus. EMBO Rep 21, e47963 (2020).

23. Faure-Dupuy, S. et al. ARL5b inhibits human rhinovirus 16 propagation and impairs macrophage-mediated bacterial clearance. EMBO Rep 25, 1156–1175 (2024).

24. Faure-Dupuy, S., Depierre, M., Fremont-Debaene, Z., Herit, F. & Niedergang, F. Human rhinovirus 16 induces an ICAM-1-PKR-ATF2 axis to modulate macrophage functions. J Virol 98, e01499–24.

25. Jubrail, J. et al. HRV16 Impairs Macrophages Cytokine Response to a Secondary Bacterial Trigger. Front Immunol 9, 2908 (2018).

26. Cox, J. et al. Accurate Proteome-wide Label-free Quantification by Delayed Normalization and Maximal Peptide Ratio Extraction, Termed MaxLFQ. Mol Cell Proteomics 13, 2513–2526 (2014).

27. Kennani, S. E. et al. MS-HistoneDB, a manually curated resource for proteomic analysis of human and mouse histones. Epigenetics and Chromatin 10, undefined-undefined (2017).

28. Tuthill, T. J. et al. Mouse respiratory epithelial cells support efficient replication of human rhinovirus. J Gen Virol 84, 2829–2836 (2003).

29. Warner, S. M., Wiehler, S., Michi, A. N. & Proud, D. Rhinovirus replication and innate immunity in highly differentiated human airway epithelial cells. Respiratory Research 20, 150 (2019).

30. Dautry-Varsat, A., Ciechanover, A. & Lodish, H. F. pH and the recycling of transferrin during receptor-mediated endocytosis. Proc Natl Acad Sci U S A 80, 2258–2262 (1983).

31. Valdés-Mora, F. et al. Acetylation of H2A.Z is a key epigenetic modification associated with gene deregulation and epigenetic remodeling in cancer. Genome Research 22, 307 (2012).

32. Janas, J. A. et al. Tip60-mediated H2A.Z acetylation promotes neuronal fate specification and bivalent gene activation. Mol Cell 82, 4627–4646.e14 (2022).

33. Mousnier, A. et al. Human Rhinovirus 16 Causes Golgi Apparatus Fragmentation without Blocking Protein Secretion. Journal of Virology 88, 11671–11685 (2014).

34. Doedens, J. R. & Kirkegaard, K. Inhibition of cellular protein secretion by poliovirus proteins 2B and 3A. EMBO J 14, 894–907 (1995).

35. Dodd, D. A., Giddings, T. H. & Kirkegaard, K. Poliovirus 3A Protein Limits Interleukin-6 (IL-6), IL-8, and Beta Interferon Secretion during Viral Infection. J Virol 75, 8158–8165 (2001).

36. Cornell, C. T., Kiosses, W. B., Harkins, S. & Whitton, J. L. Inhibition of Protein Trafficking by Coxsackievirus B3: Multiple Viral Proteins Target a Single Organelle. J Virol 80, 6637–6647 (2006).

37. Mostowy, S. & Shenoy, A. R. The cytoskeleton in cell-autonomous immunity: structural determinants of host defence. Nat Rev Immunol 15, 559–573 (2015).

