## Supplementary figures and images for "Human rhinovirus 16 impairs macrophage cytokine secretion by disrupting NF-κB nuclear translocation and intracellular cytokine trafficking"

### Supplementary Figure 1

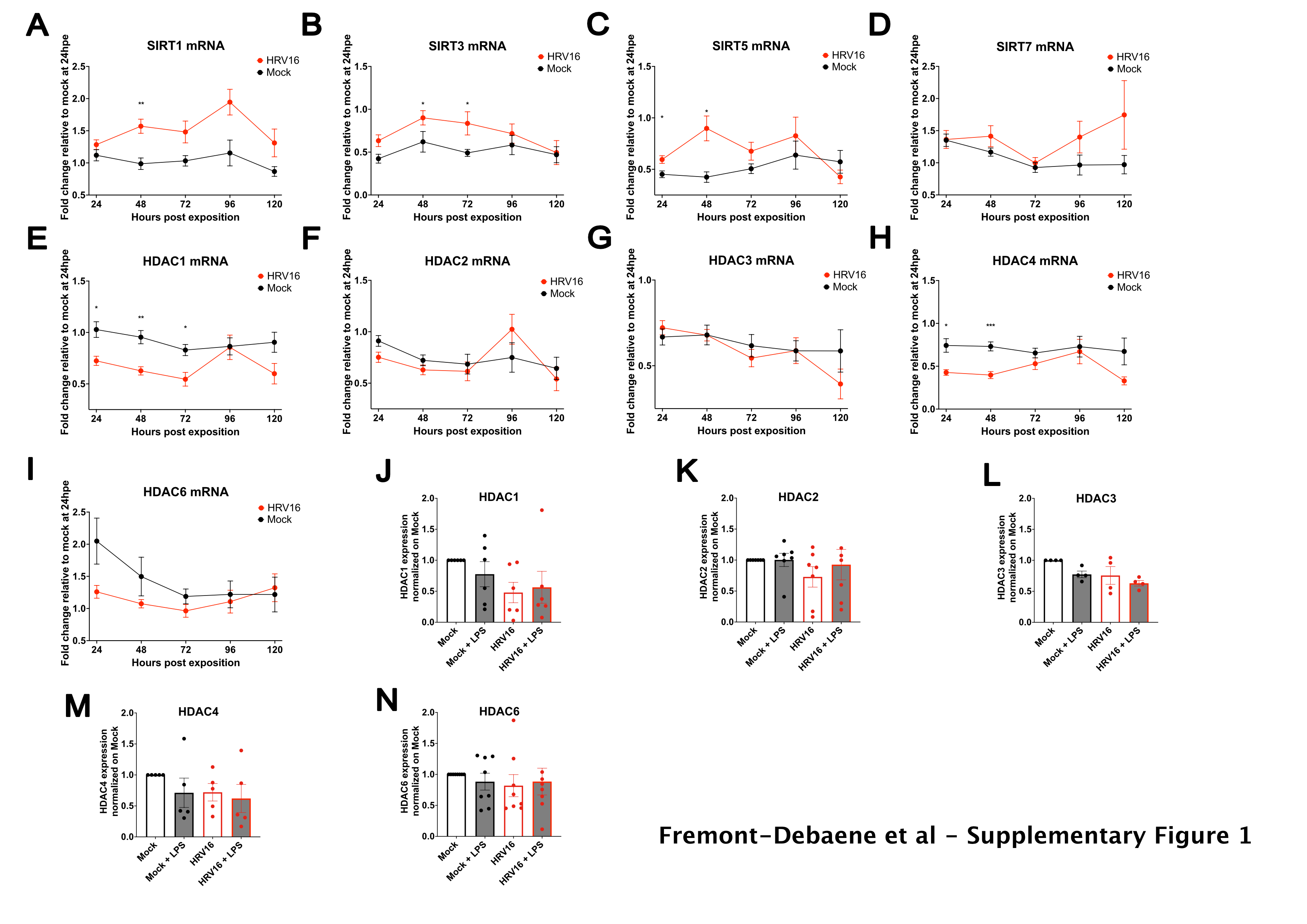
